# Patterns of microbial load and community assembly in leaf microbiomes of summer and overwintering crops

**DOI:** 10.64898/2026.05.16.725636

**Authors:** Anahi Cantoran, Peter G. Kennedy, Jannell V. Bazurto

**Affiliations:** Department of Plant and Microbial Biology, University of Minnesota, St. Paul, MN, U.S.; Biotechnology Institute, University of Minnesota, St. Paul, MN, U.S

**Author notes:** Corresponding author: Anahi Cantoran.

**Keywords:** Host-associated PCR, phyllosphere, microbial load, corn, soybean, pennycress, methylobacterium

## Abstract

Phyllosphere microbiomes are increasingly recognized as key regulators of plant health and stress responses, although they are also known to change considerably over both space and time. In the phyllosphere, members of the genus *Methylobacterium* are often abundant and ecologically important as plant growth promoting bacteria. However, knowledge about the temporal abundances and community dynamics of *Methylobacterium* in agricultural systems remains limited. To address this gap, we characterized seasonal shifts in *Methylobacterium*-specific and total phyllosphere bacterial loads and community structure on two common summer crops and one overwintering cover crop. Leaf samples of *Zea mays* (corn), *Glycine max* (soybean), and *Thlaspi arvense L*. (pennycress) plants were collected over one year in Minnesota, USA and analyzed with host-associated microbial PCR (hamPCR). Microbial loads and community composition varied strongly among hosts and across growing seasons. Corn supported the highest *Methylobacterium* and total bacterial loads, increasing towards senescence, while pennycress exhibited the lowest loads and the most distinct communities. While there were strong host-specific patterns, a group of most abundant genera were shared across all crops (*Methylobacterium*, *Sphingomonas*, *Pseudomonas*, and *Massilia)* and the most abundant *Methylobacterium* amplicon sequence variants were present on all three hosts. Our findings highlight how microbial loads and community composition change during phyllosphere assembly across diverse summer and overwintering crops, with a small core of versatile taxa dominating multiple agricultural hosts. Understanding these host and season-linked patterns provides a foundation of harnessing *Methylobacterium* strains to enhance crop productivity and resilience.

## 1. INTRODUCTION

Microbial communities associated with plants are increasingly recognized as critical components of terrestrial ecosystems, influencing host physiology, stress resilience, and biogeochemical cycling (Schimel & Schaeffer, 2012; Vandenkoornhuyse et al., 2015; Zamioudis & Pieterse, 2012). While there has been considerable research on plant-associated microbiomes belowground, aboveground leaf surfaces (i.e., the phyllosphere) also represent dynamic microbial habitats that modulate plant-environment interactions (Delmotte et al., 2009). Phyllosphere bacteria in particular have been shown to provide defense against a wide range of pathogens (Ritpitakphong et al., 2016; Vogel et al., 2021) as well as mediate responses to abiotic stressors such as drought and ultraviolet light (Lin et al., 2023; Wang et al., 2023). As such, understanding the composition, seasonal dynamics, and functional potential of phyllosphere bacterial communities is an important foundation for predicting plant responses to environmental change.

Among phyllosphere-associated bacterial taxa, members of the genus *Methylobacterium* are often found in high abundance due to leaf emissions of volatile organic compounds (VOCs) such as methanol (Abanda-Nkpwatt et al., 2006). These bacteria facultatively consume one-carbon substrates for energy and growth, in return, providing their plant hosts with a range of benefits, including phytohormone production (Madhaiyan et al., 2007), increased abiotic stress resilience (Egamberdieva et al., 2015; Jorge et al., 2019), and protection against pathogens (Ardanov et al., 2012; Kumar et al., 2015). Despite the high abundance of *Methylobacterium* on a diverse range of plant species (Corpe & Rheem, 1989), knowledge regarding their seasonal dynamics within agricultural ecosystems remains less well-documented. Recent work by Leducq et al. (2022) demonstrated that *Methylobacterium* populations can undergo pronounced seasonal shifts in composition and growth strategy in the phyllosphere of different trees in eastern North America, suggesting that temporal dynamics are critical for understanding their ecological roles.

The influence of plant host on the composition of the phyllosphere microbiome has been widely recognized in both natural (Li et al., 2022) and agricultural (Bechtold et al., 2024) systems, as well as at multiple phylogenetic scales, ranging from cultivars of the same species (Wagner et al., 2016) to distantly related plants (Kembel et al., 2014). Host species as strong drivers of *Methylobacterium* community composition have also been found in previous studies (Knief et al., 2010; Mizuno et al., 2012). Environmental factors such as temperature and water, both of which vary seasonally, also play an important role in structuring the composition of the phyllosphere microbiome due to their direct impacts on both plant and microbial growth (van der Voort et al., 2016; Xu et al., 2018). Additionally, temporal changes in phyllosphere microbial composition may reflect fluctuating resource availability as leaves mature, which for *Methylobacterium*, may be closely related to methanol production. For example, Mozaffar et al. (2018) found that in maize, leaf methanol production was high during cell wall synthesis (young leaves) and degradation (senescence), suggesting strongly seasonal patterning of VOC emissions. Collectively, these findings underscore the need to track *Methylobacterium* and total microbiome community dynamics across different hosts over time to understand microbiome assembly.

Traditional approaches to studying plant-associated microbiomes, such as amplicon sequencing of marker genes, provide valuable insight into community composition but measure only the relative abundances of taxa (Props et al., 2017). As a result, critical ecological processes, such as changes in absolute abundances of microbes, hereafter referred to as microbial load, during plant development or over environmental gradients are often overlooked. Measuring microbial load is particularly important in plants, where colonization intensity and community composition may shift independently (Friedman & Alm, 2012; Yang & Chen, 2022). Another common problem with sequencing the prokaryotic microbial community on a eukaryotic host is contamination with eukaryotic sequence reads. It is estimated that 16S ribosomal RNA gene sequences originating from the host’s genome, plastids, or mitochondria can make up the majority (>80%) of sequences obtained (Lundberg et al., 2012), which has led to the development of discriminating primers (Beckers et al., 2016) or peptide nucleic acid (PNA) PCR clamps to reduce plant host amplification (Lundberg et al., 2013). However, such methods can introduce unintended bias against microbial 16S sequence reads, leading to challenges in accurately quantifying microbial abundance (Thomas et al., 2020).

Host-Associated Microbe PCR (hamPCR) offers a promising method for simultaneously estimating microbial load and taxonomic profiles by co-amplifying host and microbial DNA within the same sequencing library (Lundberg et al., 2021). By amplifying the whole microbial community, this method allows for estimates of total and lineage-specific estimates of microbial load, using plant reads to create a ratio of microbe-to-plant abundance. Regalado et al. (2020) demonstrated that phyllosphere microbial loads on wild *Arabidopsis thaliana* plants in Europe varied widely among individuals, and that some with high load had structurally simple microbial community structure while others with low load had more complex microbial community structure. Working with the same dataset, Lundberg et al., (2022) further determined that loads of two dominant genera, *Sphingomonas* and *Pseudomonas*, had notably contrasting patterns, with loads of the former being relatively similar across samples and the latter being highly variable. HamPCR has also been applied in agricultural systems, where Lundberg et al., (2021) assessed microbial loads on mature corn leaves and showed that *Sphingomonas* was often the dominant bacterial genus on that host. Collectively, these studies demonstrate that hamPCR can reveal valuable ecological insights regarding phyllosphere microbiome load and structure, but to our knowledge, this method has not been used to track microbial load changes over time or across multiple host species simultaneously.

In this study, we used hamPCR to quantify the dynamics of phyllosphere microbial load and community composition for three crop species over a one-year period. Specifically, we sampled leaves from corn (*Zea mays*), soybean (*Glycine max*), and pennycress (*Thlaspi arvense L.*) at multiple time points in the midwestern United States (U.S.). In the U.S., corn and soybean are among the dominant summer cash crops, with the U.S. producing 377 million metric tons of corn (31% of the global supply) and 169 million metric tons of soybean (28% of the global supply) in 2024 (USDA, 2025). As both corn and soybean are predominantly summer crops, there is a temporal gap in field cover and productivity during the fall and spring seasons, which has raised interest in using winter cover crops to maintain and improve soil quality. Specifically, in the midwestern U.S., field pennycress has been identified as a promising cover crop candidate, which can be seeded anytime in the fall and harvested for seed in May-June (Johnson et al., 2015). Specifically, pennycress has shown promise in suppressing weeds and increasing the seed yield of summer crops when it’s a part of a rotation system (Hoerning et al., 2020; Johnson et al., 2015), but to date there have been no studies profiling the phyllosphere microbial communities on this crop species.

Our objectives were to assess seasonal shifts in *Methylobacterium*-specific and total phyllosphere microbial load and community composition across two summer and one overwintering crop species. We tested four hypotheses: (H1) *Methylobacterium*-specific and total bacterial loads would vary across hosts. In both cases, we expected greater similarity in microbial load between corn and soybean, which complete their life cycle within a single summer growing season, relative to pennycress, which overwinters and thus experiences additional environmental filtering during winter months; (H2) *Methylobacterium*-specific and total bacterial loads would vary over time. For *Methylobacterium,* we predicted that load would be the highest late in the growing season when leaf VOC production is elevated (Gomez et al., 2021; Mozaffar et al., 2018; Portillo-Estrada et al., 2020), and that total bacterial loads would also increase steadily across the growing season and peak at leaf senescence; (H3) *Methylobacterium*-specific and total bacterial diversity and composition would change across the growing season and among hosts. Specifically, we expected phyllosphere diversity on all hosts to be the greatest during the early stages of growth but become less diverse over time as leaves reached senescence. Further, we expected that both *Methylobacterium*-specific and total bacterial communities to change in composition over time, with composition of the pennycress community being the most distinct and dynamic due to its persistence across the fall and spring growing seasons; (H4) Given the presence of *Methylobacterium* on diverse plant species, consistent with a generalist lifestyle, we hypothesized that *Methylobacterium* amplicon sequence variants (ASVs) would be closely related regardless of host. By integrating microbial load quantification with community profiling across diverse crops with different life histories, this study provides new insights into the seasonal assembly and ecological dynamics of crop-associated phyllosphere microbiomes.

## 2. METHODS

### 2.1 Study site and sample collection

Leaf samples from individuals of three crop species, corn (*Zea mays*, B37), soybean (*Glycine max,* MN0404CN), and pennycress (*Thlaspi arvense L.*, SP32) were collected in the agricultural field plots of the University of Minnesota St. Paul campus (44.994884, −93.172868). Whole corn and soybean leaves were sampled weekly from July through September of 2022 (14 and 11 weeks, respectively). Whole pennycress leaves were sampled weekly from October to November of 2022 before the first snowfall (3 weeks) and weekly in April through May (7 weeks) of 2023 (Supplemental Figure 1). To capture the maturation of plants over time, we collected the third or fourth true leaf (after the cotyledons) from 5 random plants at each sampling time point. Two leaf samples from individual plants were taken back to the laboratory to be used for bacterial isolations and/or stored at −80°C for hamPCR.

### 2.2 Bacterial isolation and identification

Endophytic and epiphytic *Methylobacterium* strains of each crop species were isolated by washing a separate whole leaf that was collected in phosphate buffered saline (PBS) to remove soil residue, followed by cutting a ∼2.5cm x 2.5cm segment of leaf and homogenizing it in PBS with a mortar and pestle, and plating serial dilutions to selective *Methylobacterium* PIPES (MP) (Delaney et al., 2013) medium supplemented with 125 mM methanol. Epiphytic *Methylobacterium* were isolated by taking a ∼2.5cm x 2.5cm segment of the same leaf sample and pressing the leaf on methanol MP medium. Individual strains were isolated by restreaking for individual colonies of *Methylobacterium* from each plant sample; three biological replicates of each isolate were stored at −80°C in 15% DMSO for further processing.

To obtain DNA from bacterial cells, we randomly selected 30 isolates of each host and streaked them onto methanol MP agar plates and incubated them at 30°C until visible individual colonies formed (3-5 days). An individual colony from each isolate was picked and added to 20 µL of nuclease free water (NFW) and then heated at 98°C for 15 minutes to denature cells. These samples were saved at −20°C and used as a source of template DNA for PCR was performed using 12.5 µL of GoTaq Green Master Mix 2X, 0.625 µL of universal bacterial primer pair F27 and 1492R (Frank et al., 2008; each at 10 µM), and 1 µL of DNA. Thermocycler settings consisted of an initial denaturing step of 2 min at 95°C, followed by 30 cycles of 95°C for 30 sec, 55°C for 30 sec, 72°C for 1 min, and a final extension of 72°C for 5 min. Successful PCRs were verified via gel electrophoresis and further cleaned up using the Promega’s PCR Clean-Up System Quick Protocol. 10 µL of cleaned PCR products were then mixed with 5 µL of the forward primer (F27) and used for Sanger sequencing at GENEWIZ (Azenta Life Sciences, NJ, U.S.). Sanger sequences for the *Methylobacterium* sequences used as phylogenetic references were deposited in the NCBI GenBank (accession numbers PX635357, PX635358, and PX635359).

### 2.3 Host-associated microbe PCR (hamPCR)

A total of 50 (± 1) mg of leaf tissue from each plant host was weighed from the collected whole leaves and put into sterile 1.5 mL microcentrifuge tubes. Plant and microbial DNA was extracted using the DNeasy Plant Pro kit (Qiagen) and the manufacturer’s instructions, with the exception of exchanging the provided bead for three 2.3 mm chrome stainless steel beads to better homogenize the samples during bead beating. Samples were bead beaten for 2 min using a MiniBeadbeater-96 (BioSpec). Final DNA concentrations were measured with Qubit and DNA quality measured with a Nanodrop.

HamPCR (Lundberg et al., 2021) was performed on the samples, targeting the *GIGANTEA* (GI) gene for each host. *GIGANTEA* is a plant specific protein that is involved in a diverse number of physiological responses such as the circadian rhythm, light regulation, flowering time, and more (Mishra & Panigrahi, 2015). While the primers developed in Lundberg et al. (2021) for corn targeted the *LUMINIDEPENDENS* (*LD*) gene, we decided to focus on the same gene, GI, for all three hosts. The decision to use GI was determined based on the difficulties we encountered with having the bacterial forward 16S rRNA primer binding to the corn sequences and resulting in poor taxonomic assignment when run on a 2 x 300bp Illumina NextSeq P1. Specific GI primers were developed for each host at the University of Minnesota Genomics Center (UMGC), targeting conserved oligonucleotides in the GI gene (and across GI alleles for corn and pennycress). The V4 region of the bacterial 16S rRNA gene was amplified with primer pair 515F-799R, alongside peptide nucleic acid (PNA) clamps and adapter sequences from Lundberg et al. (2021) to reduce host chloroplast and mitochondria amplification. The developed host sequence primers and PNA clamp sequences used can be found in Supplemental Table 1.

The hamPCR initiated with a short round (5 cycles) of PCR to tag host and bacteria sequences and prevent amplification bias. This first PCR was done with a master mix of 12.5 µL Kapa Hifi HotStart Ready mix 2X (Roche), 3.75 µL of mixed PNAs (pPNA and mPNA, each at 5 µM, Lundberg et al., 2013), 2.5 µL of host and bacteria primers mixed (0.625 µM each), 5 µL host DNA (5 ng/µL), and 1.25 µL NFW. The first PCR step consisted of an initial denaturing step of 95°C for 3 min, followed by 5 cycles of 98°C for 20 sec, 78°C for 10 sec, 58°C for 15 sec, 55°C for 15 sec, and a final extension of 72°C for 1 min. After this, the PCR products were cleaned with SPRIselect beads (0.5X (25 µL beads) followed by 0.9X (20 µL beads)). A second round of PCR (exponential reaction) was then performed with a master mix of 12.5 µL Kapa Hifi HotStart Ready mix 2X, 3.75 µL of mixed PNAs (5 µM each), 2.5 µL of universal oligo mix (2 µM), 6.25 µL of the first round of PCR product. This second round consisted of an initial denaturing step of 95°C for 3 min, followed by 25 cycles of 98°C for 20 sec, 78°C for 10 sec, 60°C for 30 sec, and a final extension of 72°C for 1 min. A final PCR clean-up was performed with SPRIselect beads and then all samples were pooled and run on a 2 x 300bp Illumina NextSeq P1 at the UMCG. All forward and reverse .fastq files are deposited in the NCBI Short Read Archive (BioProject number PRJNA1377205).

### 2.4 Bioinformatics

All raw sequences were processed with Cutadapt (version 5.0, Martin, 2011) for NextEra adapter and primer removal. DADA2 (version 1.32, Callahan et al., 2016) and R (version 4.4.0, R Core Team, 2025) were used for further processing of reads following best practices as described in the DADA2 documentation for sequence quality, error correction, paired amplicon merging, chimeric read detection, and taxonomic identification using the DADA2 version of the Silva database (version 138.2, Quast et al., 2013; Yilmaz et al., 2014). ASVs were further taxonomically identified using NCBI_BLAST+ (version 2.13.0, Camacho et al., 2009) BLASTN using the core_nt database (Sayers et al., 2025).

### 2.5 Statistical analyses

All statistical analyses and data visualization were conducted in R version 4.2.3 (R Core Team, 2024). To estimate microbial loads across hosts, the total amount of bacterial counts was summed for each sample and divided by the total amount of respective host counts from the same sample. The packages “phyloseq” and “vegan” were used to calculate alpha and beta diversity metrics (McMurdie & Holmes, 2013, Oksanen et al., 2025). Specifically, we calculated the Shannon diversity index, evenness, and richness for all hosts, and the Shannon diversity index for each host through time. These same alpha diversity metrics were run on *Methylobacterium*-only data, which consisted of ASV data from each host, filtering for bacteria classified to the genus “*Methylobacterium*” and *“Methylobacterium-Methylorubrum*” (Alleman et al., 2025; Leducq et al., 2022). Each temporal trend involving microbial load and diversity was fitted with linear, quadratic, square root, and exponential models and the best line fit was determined based on the lowest Akaike Information Criterion (AIC) score. To determine if the bacterial communities across the three hosts differed significantly, we ran permutational multivariate analyses of variance (PERMANOVA) based on both Bray-Curtis and Jaccard index dissimilarities and used non-metric multidimensional scaling (NMDS) for community visualizations. PERMANOVA tests were followed with the “pairwise adonis2” function to determine which groups were significantly different (Martinez, 2020). Lastly, we ran abundance-occupancy models for each plant host to determine the most dominant or “core” taxa (Shade & Stopnisek, 2019), which resulted in 4 bacterial taxon that were shared across all plant hosts (Supplemental Figures 2 and 5). The abundance of the shared four bacterial taxa was then plotted across time and for each host.

### 2.6 Phylogenetic analysis

To determine the similarity of the top *Methylobacterium* ASVs across hosts, a phylogenetic analysis was conducted using the Molecular Evolutionary Genetics Analysis (MEGA) software (version 11.0.13, Tamura et al., 2021). The top 15 *Methylobacterium* ASVs for each host were aligned alongside representative sequences of *Methylobacterium* species for the monophyletic groups A (*M. fujisawaense*), D (*M. marchantiae*) and B (*M. extorquens*), as described in Leducq et al. (2022). The alignment was done using the MUSCLE algorithm in MEGA, tails were trimmed, resulting in 238 bp of aligned sequence. A Jukes-Cantor model with added gamma distribution (JC + G) was run on the aligned sequences with 1000 bootstraps. The model was determined based on the lowest AIC score after running a find the best DNA model (ML) function on MEGA. The phylogenetic reconstruction was visualized in Rstudio using the ggtree library (Xu et al., 2022).

## 3. RESULTS

### *Methylobacterium*-specific and total microbial load on plant hosts

The total reads of either *Methylobacterium* or all bacteria increased over time in all cases except for pennycress, while the total reads for the plant host consistently decreased over time (Figure 1; see inset plots). Correspondingly, microbial loads increased over time, although the specific *Methylobacterium* loads differed across the three hosts (Figure 1A). Corn accumulated the highest *Methylobacterium* loads, being 100x and 10000x higher than soybean and pennycress, respectively. Near the end of the growing season, *Methylobacterium* loads increased notably on corn after 12 weeks and soybean after 10 weeks. By contrast, the *Methylobacterium* loads on pennycress had a less distinguishable temporal pattern, although higher loads were somewhat more frequent in the spring (sampling weeks 4-10) than fall growing season (sampling weeks 1-3). Total bacterial loads over time on corn and soybean were similar to the temporal trends seen with *Methylobacterium*. Conversely, pennycress accumulated significantly higher (∼10000x) total bacterial loads compared to its *Methylobacterium* load (Figure 1B).

**Figure 1.**
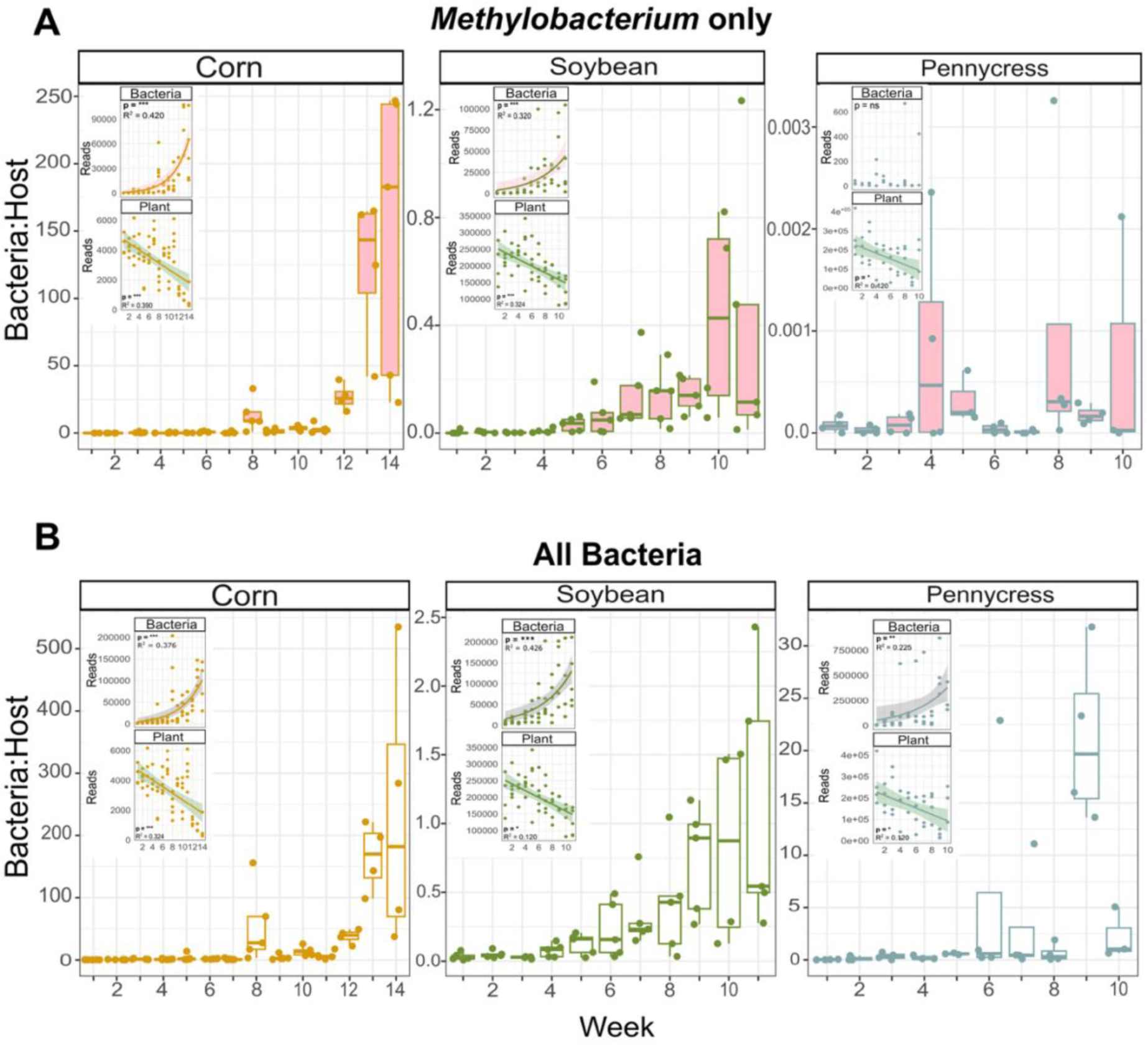
Microbial loads on the three crop species across the weeks sampled. (A) *Methylobacterium*-specific ASV loads and (B) total bacterial ASV loads. Ratios represented are as bacterial reads over plant host reads. Inset figures represent the bacteria (top) and plant (bottom) reads over time that were used to calculate the ratios for each respective facet. ANOVA p-values are presented for the reads of the bacteria and hosts, with time as the explanatory variable (ns, not significant; *, P ≤ 0.05; **, P ≤ 0.01; ***, P ≤ 0.001). Shading represents the 95% confidence interval.

### *Methylobacterium*-specific and total microbial diversity and composition on plant hosts

The diversity (Shannon H-index) of *Methylobacterium* was significantly lower on the overwintering pennycress compared to the summer grown corn and soybean (Figure 2A). Over time, *Methylobacterium* ASV diversity significantly increased for corn (p = 4 x10^-5^) and significantly decreased for soybean (p = 1.7 x10^-4^), while there was no significant temporal trend for pennycress (p = 0.7, Figure 2A). The diversity of the total bacterial ASVs was the highest on soybean, followed by corn and then pennycress (Figure 2B). In contrast to the temporal trends for *Methylobacterium* ASV diversity, the total bacterial ASV diversity significantly decreased for both corn (p = 5 x10^-3^) and soybean (p = 2 x10^-4^). No significant temporal trend was seen in the total bacterial ASV diversity for pennycress.

**Figure 2.**
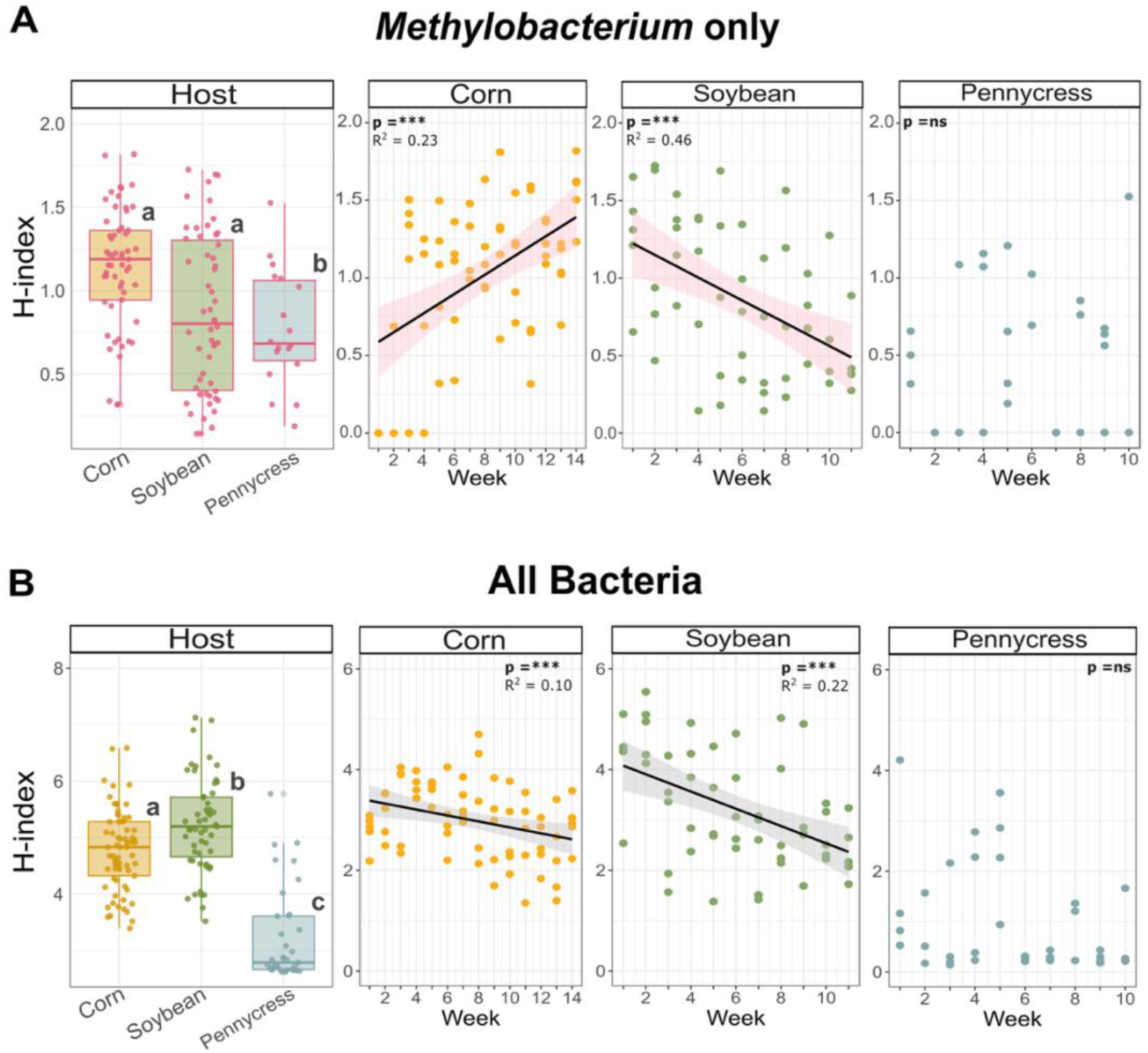
Alpha diversity across hosts and time. (A) *Methylobacterium*-specific ASV diversity across hosts and over the weeks sampled. (B) Total bacterial ASV diversity across hosts and over the weeks sampled. Significant differences among hosts based on Tukey’s HSD tests are indicated with different letters. ANOVA p-values are presented for the diversity of each host, with time as the explanatory variable (ns, not significant; *, P ≤ 0.05; **, P ≤ 0.01; ***, P ≤ 0.001). Shading represents the 95% confidence interval.

The composition of *Methylobacterium* ASVs varied across the three hosts (F_2,144_ = 8.52, p = 1 x10^-4^) (Figure 3A). Specifically, the *Methylobacterium* community on pennycress was significantly different from those on corn (p = 0.003) and soybean (p = 0.003), while corn and soybean communities were not significantly different from one another (p = 0.081). The total bacterial community composition significantly differed across the three hosts (F_2,160_ = 16.877, p = 1 x10^-4^) (Figure 3B), with all communities being significantly different from one another (each p = 0.003). The same patterns held whether the analyses were based on Bray-Curtis or Jaccard Dissimilarity (Supplemental Figure 3), indicating that differences reflected not only shifts in shared ASV abundance among hosts, but also turnover in which ASVs were present across hosts. While the overall composition of the bacterial communities on each host was distinct, four genera within the 10 most abundant bacterial genera were shared on corn, soybean, and pennycress; *Methylobacterium*, *Sphingomonas*, *Massilia*, and *Pseudomonas* (Supplemental Figure 4). Each of these genera demonstrated notable variation in abundance throughout the growing season for each host. Specifically, *Methylobacterium* and *Sphingomonas* increased in abundance for corn and soybean as plants approached the end of the growing season; *Massilia* had variable pulses of high abundance throughout the summer for corn and soybean and throughout fall and spring for pennycress; *Pseudomonas* was notably more abundant on pennycress, specifically in the fall and early spring (Figure 3C).

**Figure 3:**
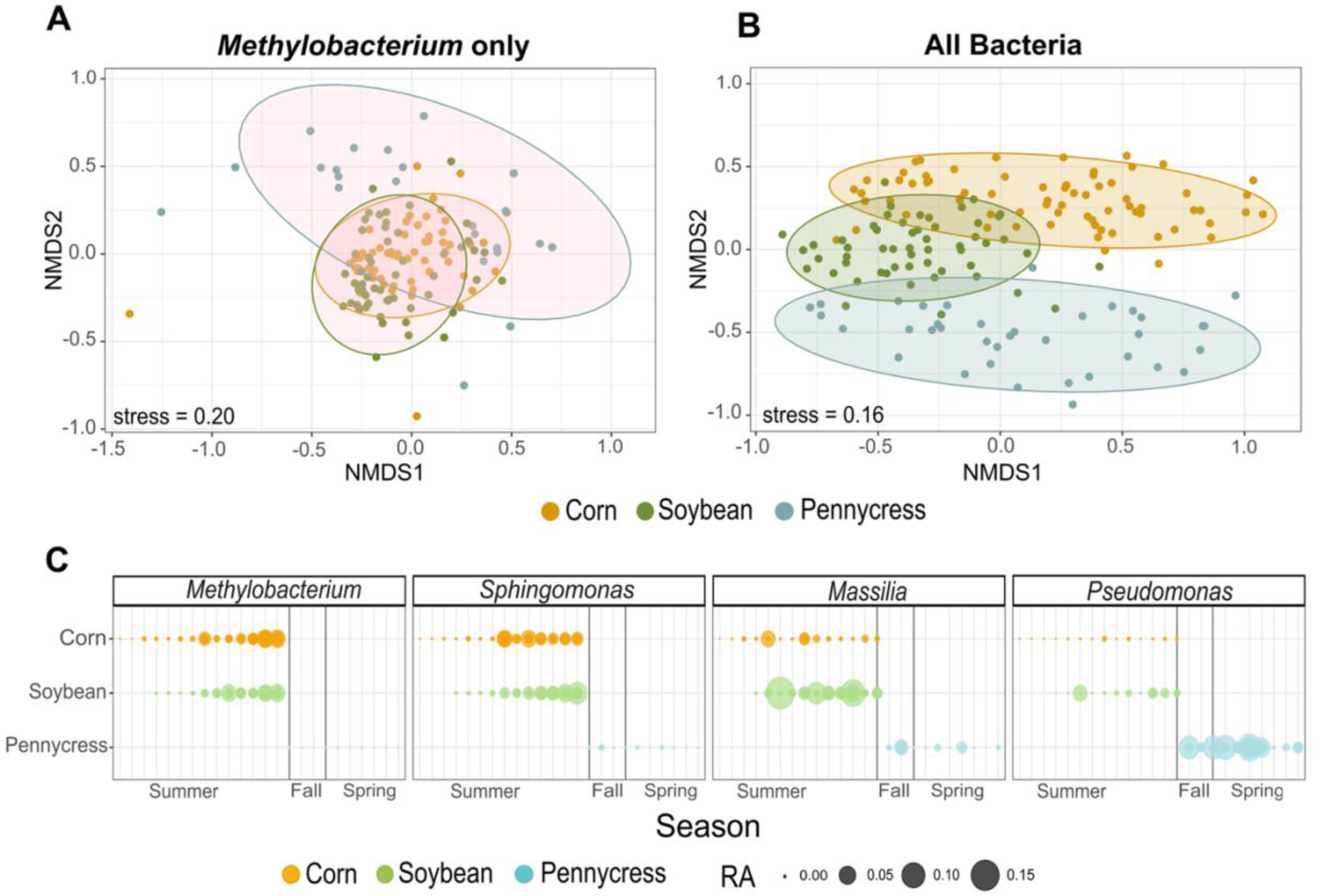
Microbial community composition across hosts. (A) NMDS of *Methylobacterium*-specific ASV composition across the three hosts. (B) NMDS of the total bacterial ASV composition across the hosts. (C) The most relative abundant (RA) bacterial genera that are shared across all three hosts across their respective growing seasons (summer for corn and soybean, fall and spring for pennycress).

Despite differences in *Methylobacterium* community composition across the three hosts, several of the most abundant *Methylobacterium* ASVs were shared. Of the top 15 most abundant *Methylobacterium* ASVs on each crop (based on total abundance over the full respective sampling periods), 7 ASVs were shared across all hosts (ASVs 1-5, 8, and 10; Figure 4). *Methylobacterium* ASV_1 was the most abundant ASV on both corn and soybean, while ASV_10 was the most abundant ASV on pennycress. The phylogenetic analysis determined that the most abundant *Methylobacterium* ASVs (including ASV_1) on corn and soybean clustered within each of the differing monophyletic groups as described in Leducq et al. (2022) (Figure 4). The most abundant *Methylobacterium* ASV on pennycress (ASV_10) clustered with *M. marchantiae*. Bootstrap values showed the greatest support for ASVs that shared similarity to *M. fujisawaense* against other groups (bootstrap value = 0.92), which then further clustered into a clade most similar to *M. extorquens* (bootstrap value = 0.97; Figure 4).

**Figure 4:**
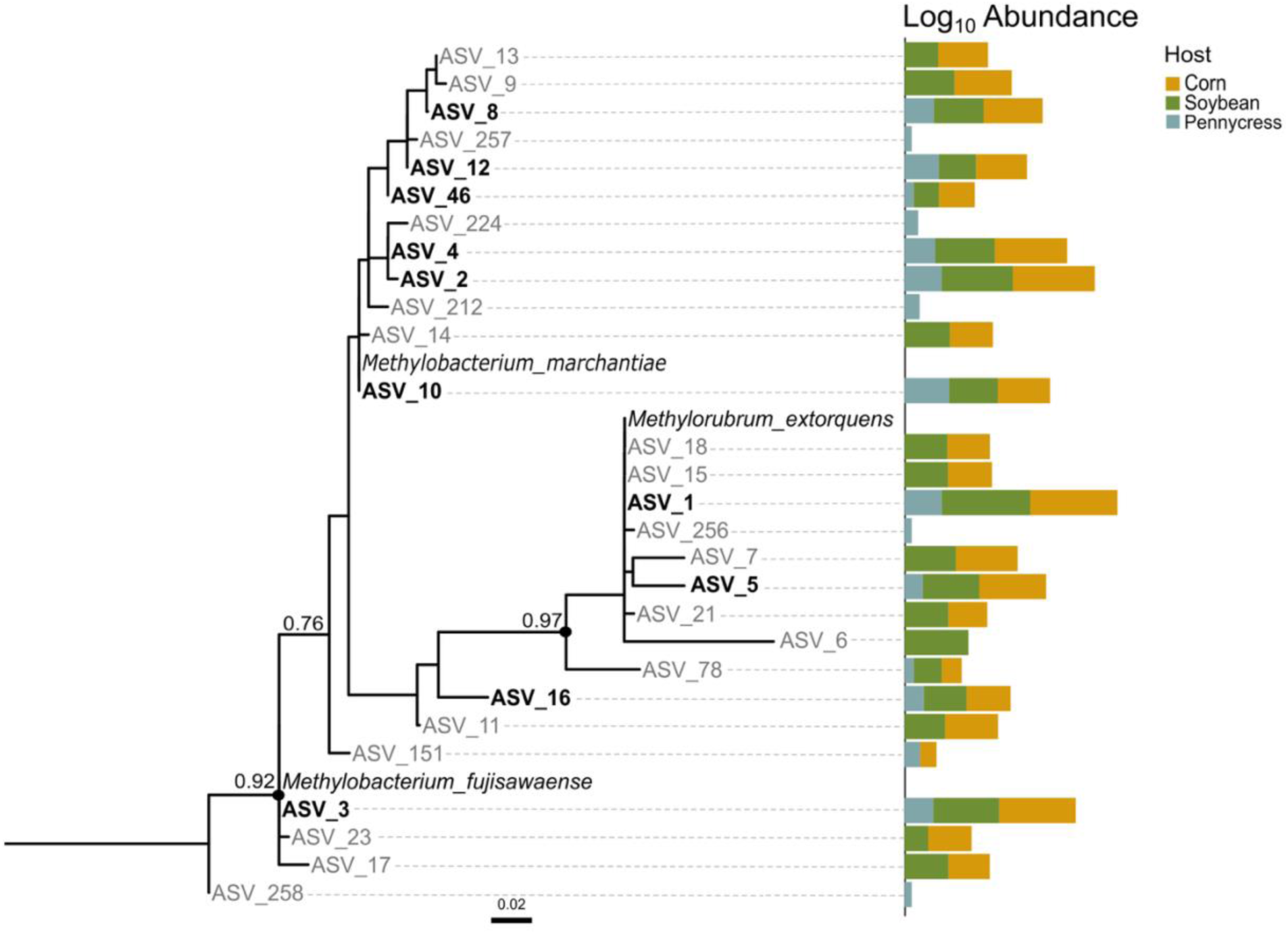
Phylogenetic comparison of the topmost abundant *Methylobacterium* ASVs per host with *M. marchantiae*, *M. extorquens*, and *M. fujisawaense* included as references. Bootstrap values below 0.7 are not presented. The total log_10_ abundance of the *Methylobacterium* ASVs are aligned with the phylogenetic tree. ASV labels in bold represent shared ASVs across all three hosts.

## 4. DISCUSSION

This study provides the first quantitative and compositional assessment of phyllosphere microbial loads across two summer and one overwintering crop species using hamPCR. In corn and soybean, microbial loads increased over the growing season, and peaked as plants approached senescence, while the overwintering pennycress exhibited less consistent temporal patterns. With regard to diversity and composition, *Methylobacterium* communities showed host-specific temporal trajectories and clear compositional differentiation among crops, while total bacterial diversity declined over time and community composition also differed significantly among hosts. Despite these differences, a set of abundant bacterial genera that were shared across hosts were observed, which consisted of *Methylobacterium*, *Sphingomonas*, *Pseudomonas*, and *Massilia*. Within *Methylobacterium*, phylogenetic analyses showed that most of the abundant ASVs were shared among the three crop species, although the most dominant ASV on pennycress belonged to a different species group than that on corn and soybean.

Our first hypothesis predicted that both *Methylobacterium*-specific and total bacterial loads would vary across hosts, with a greater similarity between the two summer annual crops, corn and soybean. This hypothesis was generally supported, as the three hosts did differ significantly in both types of loads, with the loads being significantly higher on corn. Previous culture-based work has shown that *Methylobacterium* are more readily isolated from corn relative to other crops such as cotton (McInroy & Kloepper, 1995) and soybean (Raja et al., 2008), however the magnitude of differences found in this study were much larger than we had anticipated. The substantially higher microbial loads in corn compared to the other two hosts may reflect variation in multiple factors. For example, we observed lower plant reads for corn and this may be due to plant cell wall composition alongside the DNA extraction method, which together can yield different concentrations and DNA purity (Giangacomo et al., 2021; Pipan et al., 2018). Additionally, it is important to note that DNA extractions were performed using fresh weights of plant tissue, which results in a lower dry mass as plants get older and senesce (Gland, 1963). As such, the surface area to volume that is used to achieve the same fresh weight is increased, potentially also increasing the relative amount of bacteria present. Having a lower host read count in the denominator dramatically increased microbe-to-plant read ratios for corn, although the temporal trends in both microbial and plants were similar on the two summer crops. Additionally, the total reads of *Methylobacterium* on corn and soybean were similar, suggesting both can host relatively equivalent amounts of this genus, despite appearing to have very different total microbial loads. The patterns of pernnycress, however, were quite different and we suggest other factors such as leaf surface structure, physiology, and metabolite presence, all of which can impact microbial aggregation and colonization (Omer et al., 2004; Tang et al., 2023; Vorholt, 2012; Yan et al., 2022), may underlie the differences observed. Additionally, plant hosts can have distinct antimicrobial properties that could be having an inhibitory effect on the microbial loads and composition. For example, the methanolic extracts from certain varieties of soybean can possess antibacterial properties (Hosseini Chaleshtori et al., 2017). Given that the aforementioned variables can all impact microbial composition and are not mutually exclusive, we encourage further research in understanding how microbial load (via hamPCR or other methods) is influenced by specific plant leaf properties, which ultimately impact microbial establishment and composition.

The second hypothesis predicted that microbial loads vary over time, with *Methylobacterium* peaking during periods of high leaf VOC production and total bacterial loads being higher as the end of the growing season. Previous studies have demonstrated an enrichment of *Methylobacterium* in older plants including common bean, soybean, canola, and prairie grasses (Copeland et al., 2015; Ding & Melcher, 2016), consistent with this, *Methylobacterium* loads were highest late in the growing season on both corn and soybean. The amount of methanol released is greatest, however, during the early seedling stage for plants, when plants undergo the most cell wall synthesis and pectin is demethylesterified by pectin methylesterases (Dorokhov et al., 2018; Eller et al., 2012; Mozaffar et al., 2018). Given that we delayed sampling until fully mature leaves were present, we did not sample during the methanol peaks at the seedling stage (Supplemental Figure 1). However, since there is likely a strong link between methanol production and *Methylobacterium* load, we expect that this genus would also have a high load during that phase of plant growth. Similarly, we presume the observed differences in *Methylobacterium* loads across hosts reflected different amounts of VOC emission profiles across hosts, including methanol (Kigathi et al., 2019). Further research that quantitatively measures the VOC emissions of these hosts through time in conjunction with *Methylobacterium* community characterization will clarify the relationship of VOC emissions to the loads of methylotrophs.

We further hypothesized that *Methylobacterium* and total bacterial community diversity and composition would vary across both hosts and the growing season, with the overwintering pennycress exhibiting the most distinct and dynamic phyllosphere. Across time, the total bacterial diversity decreased as plants got older across all hosts, supporting existing evidence that phyllosphere communities become more homogenous across host plants as they reach senescence (Copeland et al., 2015; Grady et al., 2019). However, this trend differed for *Methylobacterium* specific communities, where diversity patterns over time varied by host for reasons not immediately clear. In terms of community composition, our results are consistent with previous studies showing similar differences in trajectories across different hosts. For example, Copeland et al. (2015) followed the successional microbial changes in common bean, soybean, and canola across a growing season and found a higher similarity between the hosts from the family Fabaceae (common bean and soybean), which share similar physiological characteristics, than with canola (from the family Brassicaceae).

While the total bacterial community diversity and composition in the phyllosphere were generally different by both host and season, we also observed the presence of a shared ‘core ‘phyllosphere community (i.e., microbial taxa that are consistently found across time, space, and/or host) (Shade & Stopnisek, 2019), composed of *Massilia*, *Methylobacterium*, *Pseudomonas*, and *Sphingomonas*. These core bacterial genera are common dominant colonizers of the phyllosphere across other plants such as *Arabidopsis*, canola, common bean, switchgrass, and miscanthus (Almario et al., 2022; Copeland et al., 2015; Grady et al., 2019). Our study demonstrates that *Methylobacterium* is the most dominant bacterium on corn and soybean plants, yet sparse on overwintering pennycress, further emphasizing the dominant yet dynamic nature of *Methylobacterium*. We suspect that the variable load and diversity on pennycress leaves may be in part explained/constrained by the additional environmental filtering between the late fall and early spring months (e.g., ice-thaw cycles, short day length, low light, desiccation, and other), which is unfavorable for bacteria not adapted for those conditions. Interestingly, in the only other study utilizing hamPCR in an agricultural setting, *Methylobacterium* was largely absent on mature corn leaves, yet the other three dominant genera in our system (*Massilia*, *Pseudomonas*, and *Sphingomonas*) were common and abundant (Lundberg et al., 2021). Why *Methylobacterium* was missing in that system, but highly abundant in ours could be due to many factors (e.g., different corn varieties, location, different abiotic conditions and resources), suggesting future research comparing *Methylobacterium* loads across corn varieties co-planted in different geographic areas will determine the relative importance of host versus environment in determining abundances of important plant growth promoting genus.

Regarding our fourth hypothesis, given the widespread occurrence of *Methylobacterium* across plant hosts, we expected high similarity among ASVs across crops. Despite clear host effects on total bacterial composition, the most abundant *Methylobacterium* ASVs were shared across all hosts, with several ASVs closely related to *M. extorquens*. This species is a common phyllosphere inhabitant and belongs to one of three *Methylobacterium* clades frequently found on plants (Leducq et al., 2022). The overlap in ASVs between corn and soy likely reflects dispersal from shared environmental sources, as previous studies have shown that surrounding vegetation strongly influences phyllosphere community assembly (Lajoie & Kembel, 2021; Meyer et al., 2022). This interpretation is consistent with the short proximity of our corn and soybean plots (Supplemental Figure 1) and similar to a recent study by Meyer & Lindow (2025), who found that the phyllosphere of corn and soybean was similar to that of the nearest surrounding herbaceous and woody vegetation. Soil microbial communities shaped by crop rotations may also serve as an important reservoir for *Methylobacterium*, influencing aboveground assembly that can potentially affect plant performance/crop yield (Benitez et al., 2021; Yang et al., 2023). Given that the pennycress plants were, however, also co-located and grown on the same soils as those of corn and soybean, the clear shift to a different *Methylobacterium* as the dominant ASV suggests that community microbial environmental filtering may still be strongly mediated the host plant. Whether ASV_10 helped pennycress with more challenging abiotic conditions could not be assessed with our data, but this result does suggest that different crop species may have *Methylobacterium* clade specific preferences (Raja et al. 2008). Future research comparing *Methylobacterium* composition across different crop species at larger time intervals, spatial distances, and different land-use histories will help parse the major contributors to the assembly of *Methylobacterium* communities on these crops.

In conclusion, by combining microbial load quantification with community profiling using hamPCR, we have shown how abundance and compositional data can enhance our understanding of ecological processes underlying phyllosphere assembly. Our findings also build on recent phyllosphere research in Minnesota (Meyer et al., 2022; Meyer & Lindow, 2025) by revealing that *Methylobacterium* load, diversity and composition shifted markedly across seasons, with a small set of ASVs colonizing all hosts. Understanding these seasonal patterns can guide the selection and application of naturally occurring *Methylobacterium* as bioinoculants to enhance growth and resilience. Future work should focus on how host traits and environmental conditions regulate *Methylobacterium* establishment their ability to modulate VOC emissions. This exploration will be key to predicting and optimizing plant-microbe interactions in agricultural systems.

## Acknowledgements

We are grateful to the members of the Bazurto and Kennedy labs and Drs. Jennifer Powers, Trinity Hamilton, and Jonathan Schilling for reviewing drafts of this manuscript. We thank Dr. Aaron J. Lorenz, Dr. Nathan Springer, and Dr. David Marks for providing access to the field plots of each plant host; Peter Hermanson, Sonia Bolvaran, and Eric Warme for their assistance on the field; Dr. Anthony Brusa for his assistance in identifying homologous pennycress genes for this study; Lauren Popel for her help with sample collection and bacterial isolations; Mark Murphy for his help with hamPCR protocol development; Trevor Gould for his assistance with the bioinformatics.

## SUPPLEMENTAL FIGURES

**Supplemental Figure 1:**
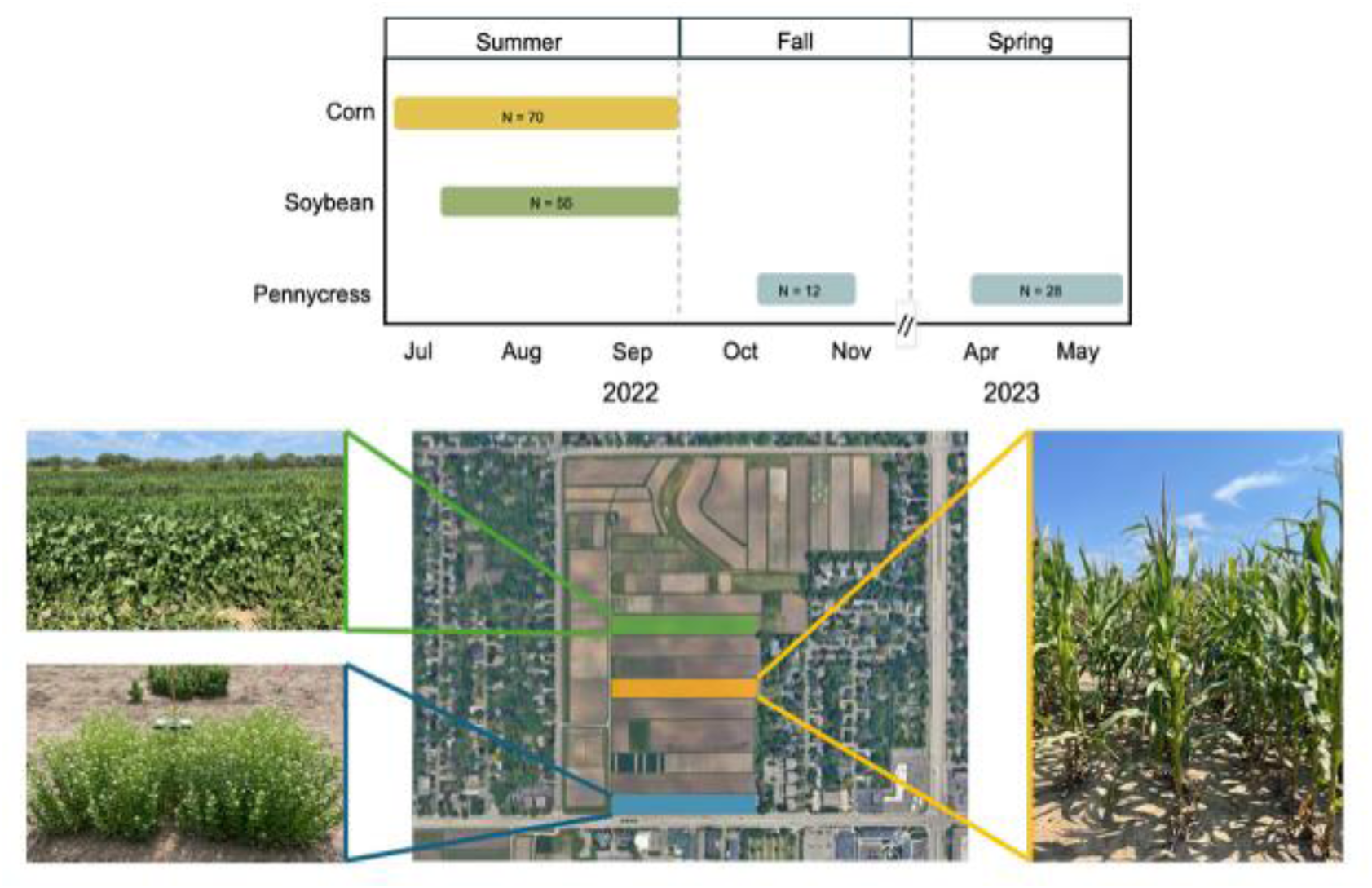
Field sampling layout. Top panel shows the time frames when samples were collected for the respective crops. Bottom panel demonstrates the agricultural field from which plants were sampled from with respective images of host plants. Soybean (top left, green plot), pennycress (bottom left, blue plot), and corn (right, yellow plot). The total agricultural field is approximately 420,000 m^2^.

**Supplemental Figure 2:**
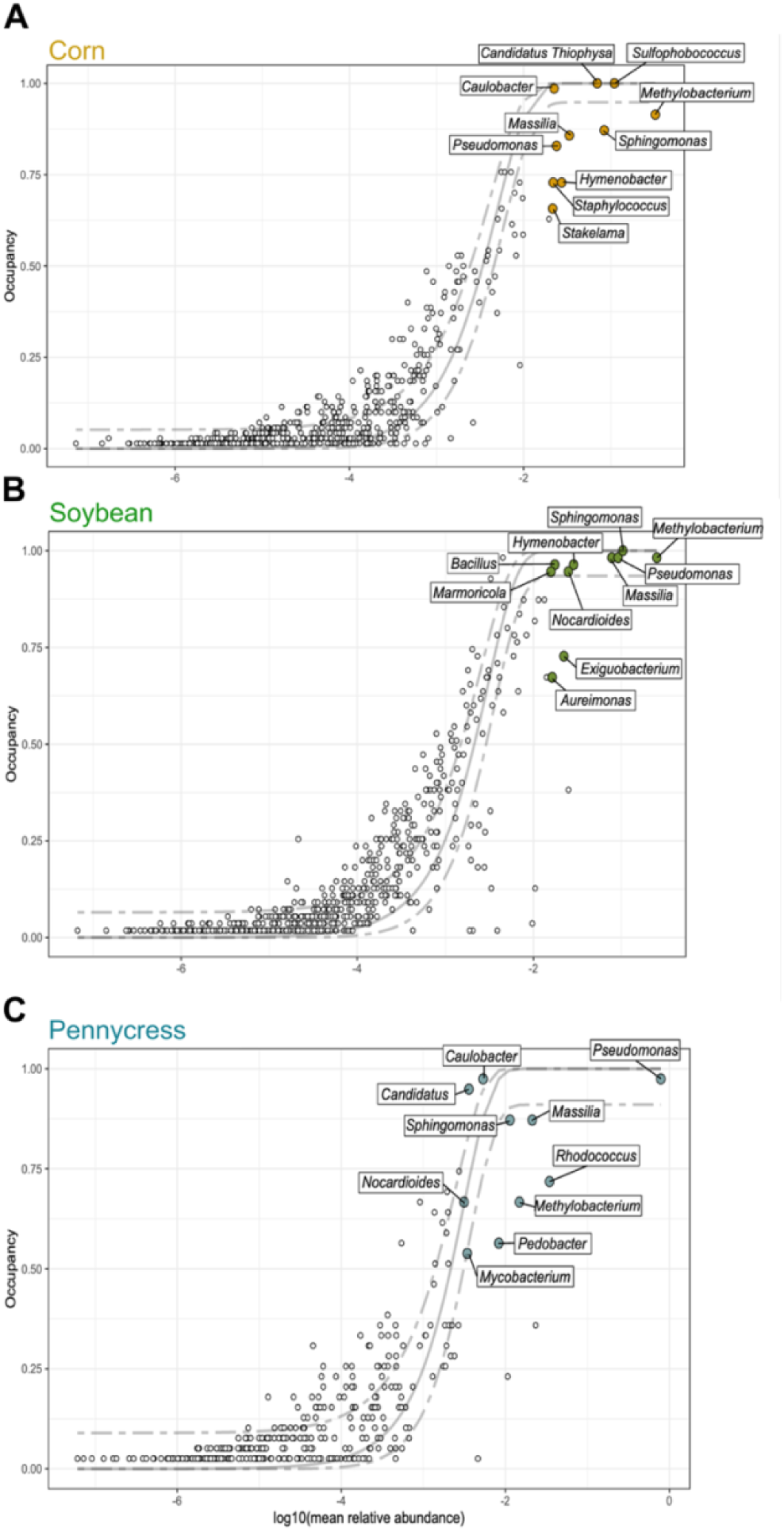
Abundance-occupancy relationships of the bacterial communities on (A) corn, (B) soybean, and (C) pennycress. Colored points indicate taxa belonging to the 95^th^ percentile of both abundance and occupancy. Points in white represent each ASV of the community. Gray solid line represents predicted abundance-occupancy neutral model with 95% confidence intervals. Adapted from Shade and Stopnisek (2019).

**Supplemental Figure 3:**
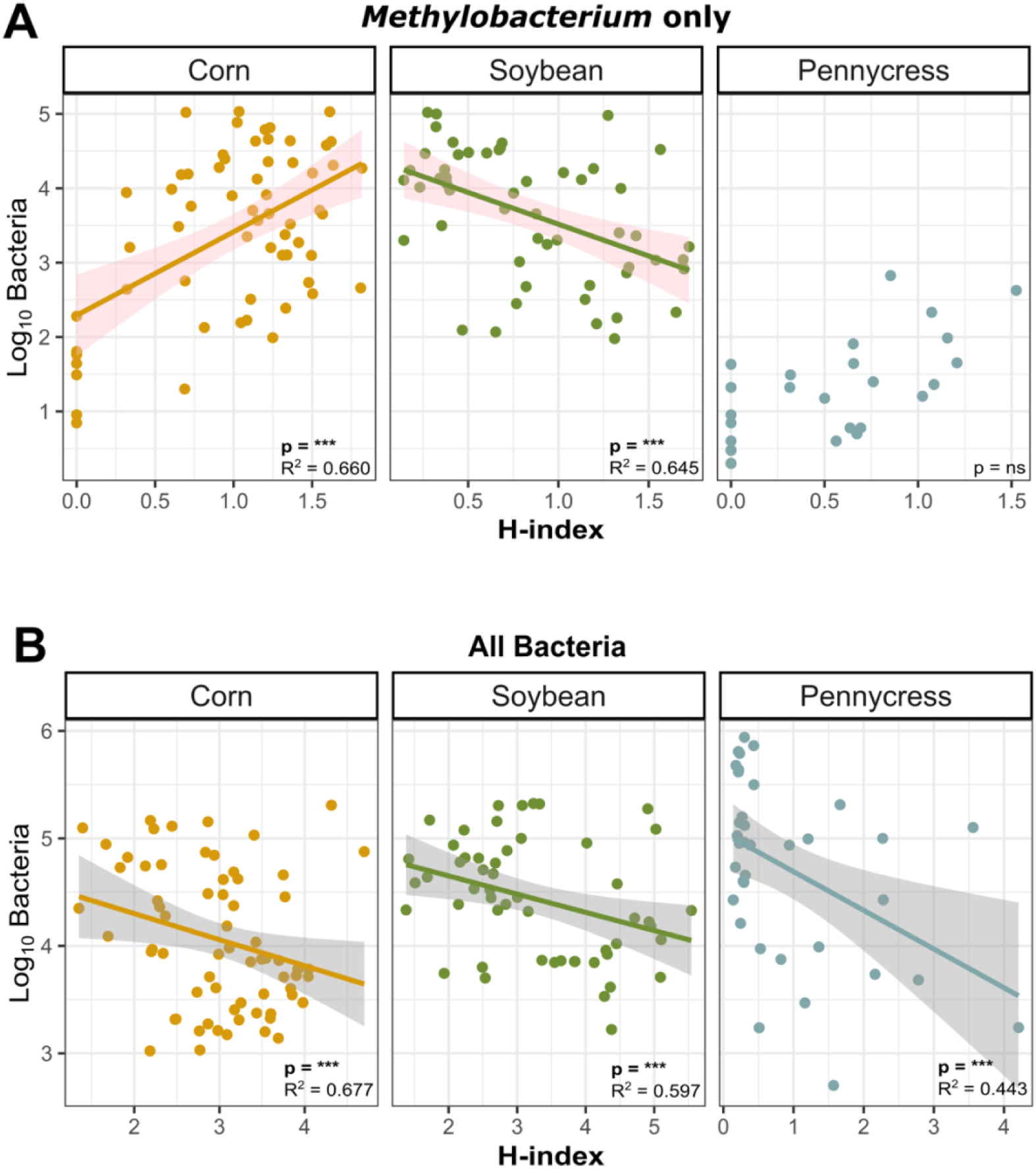
Relationship between bacterial load (Log_10_) and Shannon H-index diversity for (A) *Methylobacterium*-specific ASVs and (B) total microbial ASVs.

**Supplemental Figure 4:**
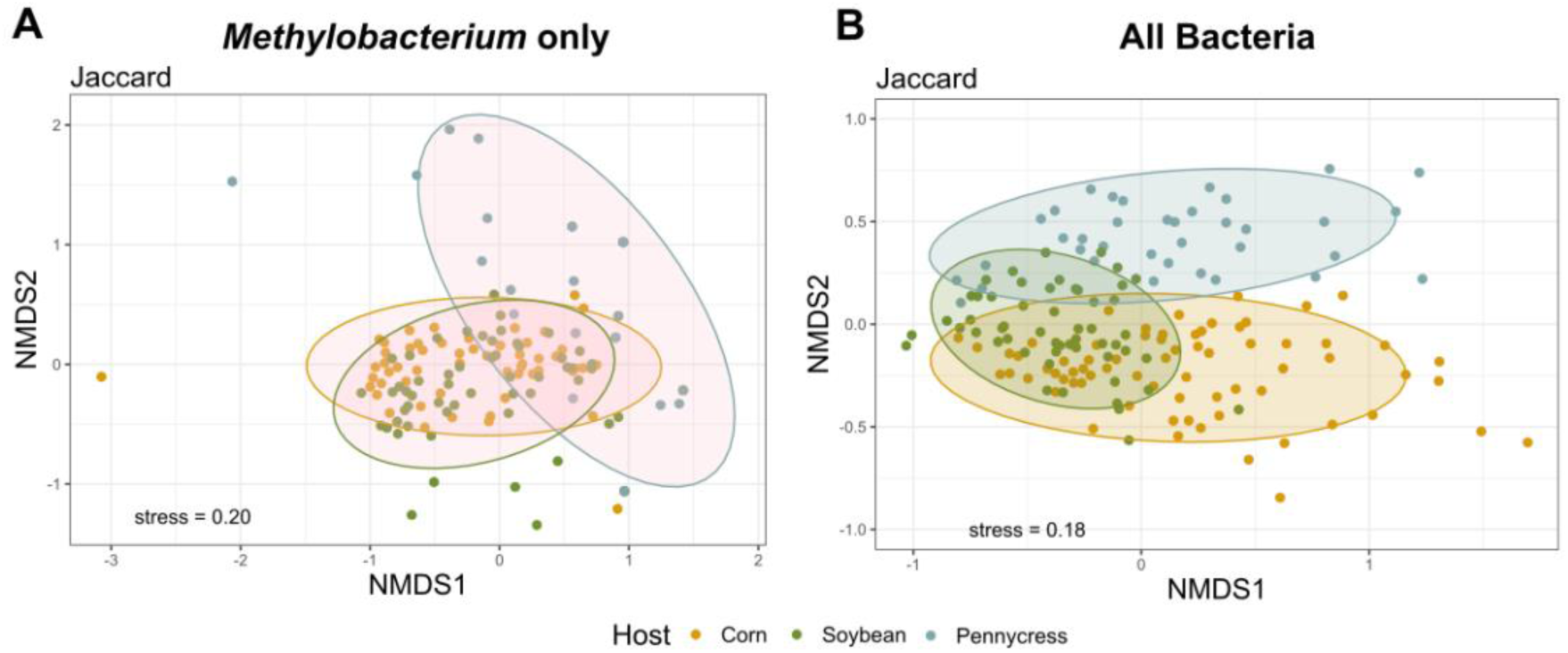
NMDS of (A) *Methylobacterium*-specific communities and (B) total bacterial communities based on Jaccard dissimilarity.

**Supplemental Figure 5:**
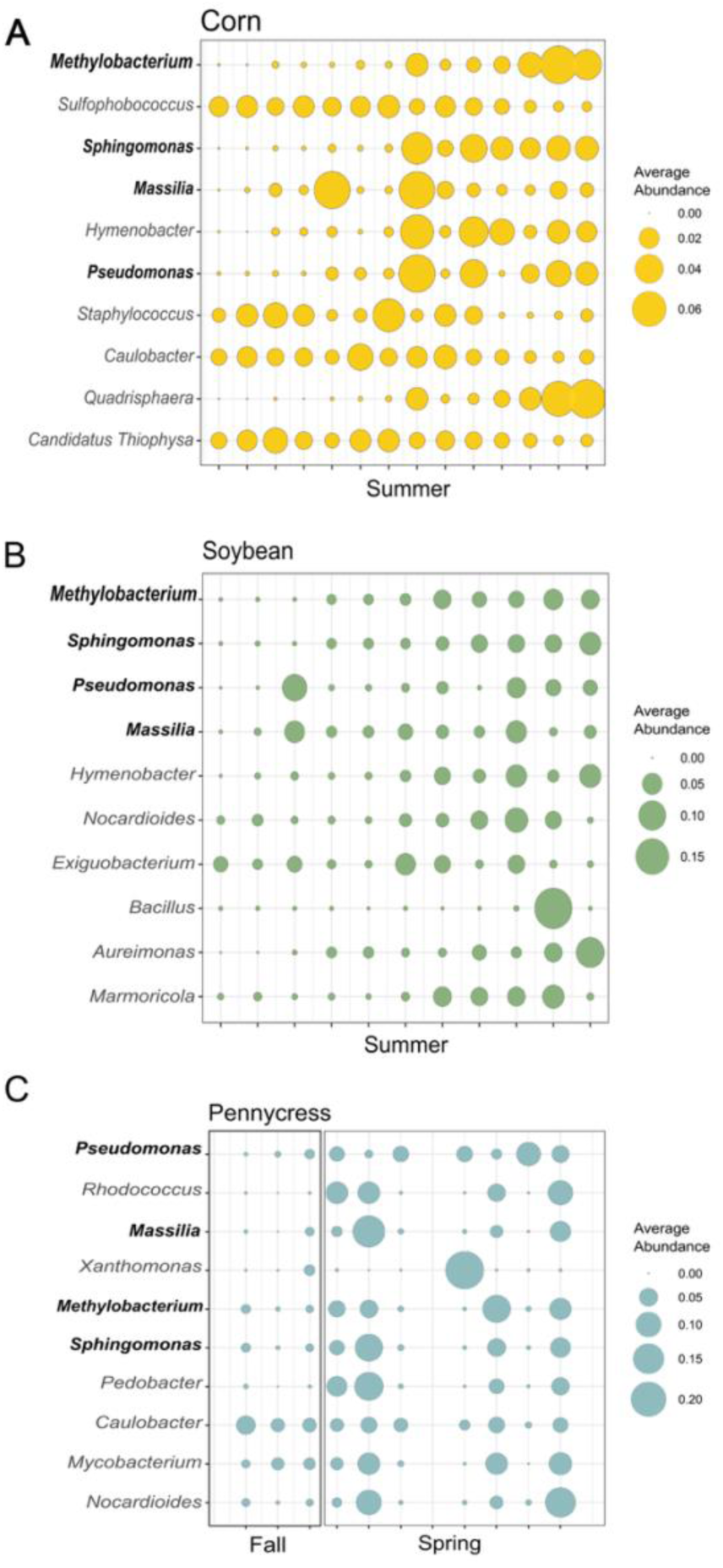
The 10 most abundant bacteria in (A) corn, (B) soybean, and (C) pennycress over time based on their total average abundance at that sample time. Values in the x-axis represent the week of the year during the indicated season.

